# Molecular subtyping improves prognostication of Stage 2 colorectal cancer

**DOI:** 10.1101/674614

**Authors:** Rachel V Purcell, Sebastian Schmeier, Yee Chen Lau, John F Pearson, Francis A Frizelle

## Abstract

Post-surgical staging is the mainstay of prognostic stratification for colorectal cancer (CRC). Here, we compare TNM staging to consensus molecular subtyping (CMS) and assess the value of subtyping in addition to stratification by TNM. Three hundred and eight treatment-naïve colorectal tumours were accessed from our institutional tissue bank. CMS was carried out using tumour gene-expression data. Staging and CMS were analysed with respect to clinicopathologic variables and patient outcome. CMS alone was not associated with survival, while TNM stage significantly explained mortality. Addition of CMS to TNM-stratified tumours showed a prognostic effect in stage 2 tumours; CMS3 tumours had a significantly lower overall survival (*P* = 0.006). Stage 2 patients with a good prognosis showed immune activation and up-regulation of tumour suppressor genes. Although stratification using CMS does not outperform TNM staging as a prognostic indicator, gene-expression based subtyping shows promise for improved prognostication in stage 2 CRC.

## Background

Colorectal cancer (CRC) is a highly heterogeneous disease with different outcomes, therapeutic response and clinical behavior, even within the same clinically designated staging and grading systems. Recent attention has been focused on the molecular mechanisms underlying CRC, and developing novel approaches to stratify tumours into subgroups that have clinical utility for improved prognostication and targeted therapy.[1]

Previous molecular classification systems have relied on combinations of molecular features, including *BRAF*, *KRAS and TP53* mutation status, microsatellite instability, CpG island methylator phenotype, somatic copy number alterations, and activation of various molecular pathways such as WNT and MYC, in order to classify CRC into subgroups.[2–5] Associations with clinical and histological features and outcome have been reported using these subtyping classification systems, and many show overlapping characteristics. However, discrepancies exist between these classification systems, and they have not been widely embraced in the clinical setting for CRC, due to technological limitations and cost of undertaking the necessary laboratory tests, as well as the question of the impact of tumour heterogeneity on results.[6, 7]

The advent of high-throughput sequencing has ushered in a new era of subtyping studies, and in 2015, the Colorectal Cancer Subtyping Consortium published a novel classification system based solely on gene expression data that drew largely on six previously published molecular classification systems.[8] The introduced classifier stratifies CRC into four consensus molecular subtypes (CMS). In particular, the value of using CMS for prognostication and precision medicine has been highlighted in several recent publications.[9] However, in the absence of targeted therapy regimens for primary CRC, the value of stratifying tumours using CMS as a prognostic tool has yet to be evaluated.

In order to assess the utility of CMS as a prognostic tool for CRC, we have evaluated molecular subtyping in relation to clinicopathologic features and patient outcome in a large single-institution cohort of treatment-naïve CRC tumours, and compared the findings to that of standard histological classification.

## Material and Methods

### Patients

Colorectal cancer (CRC) tissue samples banked at the Cancer Society Tissue Bank (University of Otago, Christchurch, New Zealand), with informed written consent were used in this study. Patient data, including staging, recurrence, metastases, treatment and histology was retrospectively collected from patient medical records. Exclusion criteria included patients with hereditary CRC, and patients who had received pre-operative chemotherapy or radiation therapy, resulting in a cohort of 308 patients. This study was carried out with ethical approval from the University of Otago Human Ethics Committee (ethics approval number: H16/037).

### RNA extraction

Tumour core samples were dissected from surgical specimens and immediately frozen in liquid nitrogen and initially stored at −80°C. RNA was extraction was carried out as detailed previously.[10] Briefly, RNA was extracted from < 20 mg of tissue using RNEasy Plus Mini Kit (Qiagen), including DNAse treatment, following tissue disruption using a Retsch Mixer Mill. Purified RNA was quantified using the NanoDrop 2000c spectrophotometer (Thermo Scientific, Asheville, NC, USA), and stored at −80°C.

### RNA sequencing

RNA-sequencing was carried out using the Illumina HiSeq 2500 V4 platform to produce 125bp paired end reads, as previously described.[10] In brief, library preparation, including ribosomal RNA depletion using RiboZero Gold, was carried out using Illumina TruSeq V2 reagents. The libraries were sequenced on 3 x 5 lanes of an Illumina HiSeq 2500 instrument.

### RNA sequencing data processing

Low quality read segments, remnant adaptor sequences and very short reads were subsequently removed using fastq-mcf from ea-utils (v 1⋅1⋅2⋅779)[11] and SolexaQA++ with default parameters (v3⋅1⋅7⋅1)[12]. Salmon (v0⋅11⋅2)[13] was used to quantify transcript expression of GRCh38 (Ensembl release 93). Gene-level tags-per-million (TPM) counts were derived using the tximport package (v1⋅6⋅0)[13]. The data processing protocols can be accessed at https://gitlab.com/schmeierlab/crc/crc-nz-2018.

### Consensus molecular subtyping

The Single Sample Predictor (SSP) method available in the CMS classifier (v1⋅0⋅0, https://www.synapse.org/#!Synapse:syn4961785) R package[8] was used to classify samples into molecular subtypes of colorectal cancer based on derived TPM values for genes. Similarity between gene expression profiles is calculated as Pearson’s correlation of log2 scaled values, and a sample is considered to be similar to a centroid if the correlation is at least 0⋅15. In order for a sample to be classified, a correlation to the most similar centroid has to be higher by 0⋅06 than a correlation to the second most similar centroid. These values are set by default in the SSP method. The data processing protocols can be accessed at https://gitlab.com/schmeierlab/crc/crc-nz-2018.

### Differential gene expression analysis

We used count tables from the RNA sequencing data processing, in addition to the class information produced through the CMS classification step, and ran a differential gene expression analysis for all genes. We used each sample within a given subtype as a replicate of that subtype, and ran the edgeR (v3⋅20⋅7)[14] package to compare each subtype against each other, extracted genes that are up- or down-regulated, using a Benjamini and Hochberg false-discovery rate[15] adjusted *P*-value (< 0⋅05), and a log2 fold-change greater or smaller than zero. Only genes were considered that were differentially expressed in all comparisons for a subtype against all other subtypes. The data processing protocols can be accessed at https://gitlab.com/schmeierlab/crc/crc-nz-2018.

### Enrichment analysis

We used the differentially expressed genes per CMS subtype, and the type of expression (up-or down-regulation) and sub-selected the top 500 genes and input to the clusterProfiler package (v3⋅6⋅0)[16] for term enrichment analysis. We sub-selected terms based on an FDR adjusted *P*-value < 0⋅1 for enrichment of the gene-set in biological categories. The biological categories and corresponding gene-sets used in the analysis were extracted from MSigDB[17] (version 6⋅1). We sub-selected the following categories for the analysis: KEGG, REACTOME, BIOCARTA, PID, HALLMARK GENES, and Gene Ontology (GO) biological processes. The data processing protocols can be accessed at https://gitlab.com/schmeierlab/crc/crc-nz-2018.

### Statistical analysis

Associations between classifiers and clinicopathological variables were assessed by Chi-square tests or Fisher’s exact test, if there were expected cell counts less than one, or if 80% of cells had counts less than five. For tables larger than two by two, *P*-values were computed with Monte-Carlo simulation. Kaplan-Meier survival curves were calculated using 5- and 10-year estimates of survival for overall survival (OS) and progression-free survival (PFS). Association of classifiers with OS and PFS progression was assessed using Cox proportional hazard models with and without clinicopathological covariates. Significance of multilevel factors was assessed with Chi-squared tests on analysis of deviance. Examination of residual plots showed that modelling assumptions were valid. All statistical tests were 2-sided and considered significant at a *P*-value of 0⋅05, and all analysis was performed in R 3⋅4⋅3 (Vienna, Austria).

## Results

### Patient cohort

Colorectal cancers from 308 patients were included (median age 73⋅7 years; range, 28–91 years). One hundred and sixty-three patients were female and 145 were male, with 296 of the patients of European decent, three Asian and nine Maori. Right-sided tumours were more common (55%) compared to left-sided colon tumours (27%) and rectal tumours made up 18%. Median follow up was 50 months (range, 0⋅3–172 months). More detailed patient demographics are shown in Supplementary Table S1.

### Association of TNM staging with clinical variables

Post-surgical TNM staging based on pathological examination stratified the cohort as follows: 53 stage 1, 128 stage 2 patients, 105 stage 3 and 22 stage 4 patients. Analysis of associations between post-surgical staging of patients and clinicopathological variables (Tables 1 and 2) showed that increasing TNM stage was significantly associated with lymph-node positivity and subsequent development of metastasis, which can be attributed to liver metastases; there was no association with local recurrence.

**Table 1.** Tumour recurrence, metastasis and lymph-node invasion by Consensus Molecular Subtype and by post-operative stage

|  | LR<br>% | Mets<br>% | Mets + LR<br>% | LM<br>% | LN<br>% |
| --- | --- | --- | --- | --- | --- |
| **CMS1** | 3.3 | 11.7 | 13.3 | 3.3 | 25 |
| **CMS2** | 4.1 | 26.2 | 26.9 | 21.4 | 37.9 |
| **CMS3** | 5.3 | 13.2 | 18.4 | 7.9 | 44.7 |
| **CMS4** | 0 | 41.2 | 41.2 | 29.4 | 64.7 |
| **P-value\*** | 0.9061 | 0.0211 | 0.0479 | 0.0009 | 0.0171 |
| **TNM1** | 0 | 11.3 | 11.3 | 9.4 | 0 |
| **TNM2** | 4.7 | 9.4 | 12.5 | 6.2 | 1.6 |
| **TNM3** | 6.7 | 31.4 | 32.4 | 20 | 97.1 |
| **TNM4** | 4.5 | 100 | 100 | 81.8 | 86.4 |
| **P-value** | 0.2407 | < 0.0001 | < 0.0001 | < 0.0001 | < 0.0001 |

**Table 2.** Histological characteristics of colorectal cancer tumours by Consensus Molecular Subtype and by post-operative stage

|  | **Poorly diff** | **Mucinous** | **LVI** | **EMVI** | **PNI** |
| --- | --- | --- | --- | --- | --- |
|  | % | % | % | % | % |
| **CMS1** | 48.3 | 20 | 31.7 | 13.3 | 8.3 |
| **CMS2** | 9 | 2.8 | 28.8 | 15.2 | 11.7 |
| **CMS3** | 10.5 | 18.4 | 31.6 | 7.9 | 7.9 |
| **CMS4** | 17.6 | 11.8 | 35.3 | 23.5 | 23.5 |
| **P-value\*** | < 0.001 | 0.0001 | 0.8686 | 0.4497 | 0.3334 |
| **TNM1** | 11.4 | 7.5 | 11.3 | 0 | 0 |
| **TNM2** | 18 | 12.5 | 23.4 | 7 | 9.4 |
| **TNM3** | 18.1 | 9.5 | 46.7 | 26.7 | 18.1 |
| **TNM4** | 22.7 | 9.1 | 72.7 | 40.9 | 31.8 |
| **P-value** | 0.5737 | 0.7864 | < 0.001 | < 0.001 | < 0.001 |

Tumour recurrence, metastasis and lymph-node invasion by Consensus Molecular Subtype (CMS) and by tumour-node-metastasis (TNM) stage given as percentages. *P*-value* is derived from analysis of classified tumours only and a *P*-value of < 0.05 is considered significant; LR, local recurrence; Mets, distant metastasis, diagnosed either at surgery or during the follow-up period; Mets + LR, local recurrence and distant metastasis combined; LM, liver metastasis; LN, lymph-node positive.

Histological characteristics of colorectal cancer tumours by Consensus Molecular Subtype (CMS) and by tumour-node-metastasis (TNM) stage given as percentages. *P*-value* is derived from analysis of classified tumours only and a *P*-value of < 0.05 is considered significant. Poorly diff, poorly differentiated; LVI, lymphovascular invasion; EMVI, extramural venous invasion; PNI, perineural invasion.

### CMS subtypes and clinical variables

Of the 308 patients, 60 were classified as CMS1 (19%), 145 as CMS2 (47%), 38 as CMS3 (12%) and 17 as CMS4 (6%) (Supplementary Table S2). Univariate analysis of patient demographic and clinical variables showed that CMS1 tumours were more likely to be right-sided, found in females, poorly-differentiated, with a high proportion of mucinous histology and less likely to be seen in younger patients. CMS2 tumours made up nearly half of our cohort and were predominantly left-sided tumours found in male patients, and showed a negative association with mucinous type. CMS4 tumours were associated with younger age, and presented at an advanced TNM stage with lymph node positivity. There was no significant difference in the local recurrence rates between subtypes, but CMS2 and CMS4 were associated with higher rates of distant metastases and this association was attributable to liver metastases. A detailed breakdown of associations with CMS subtypes is given in Tables 1, 2 and 3.

**Table 3.** Patient and tumour characteristics by Consensus Molecular Subtype

|  |  |  | **CMS1**<br>**(n=60)** | **CMS2**<br>**(n=145)** | **CMS3**<br>**(n=38)** | **CMS4**<br>**(n=17)** | ***P*-value\*** |
| --- | --- | --- | --- | --- | --- | --- | --- |
|  |  | (n) | % | % | % | % |  |
| **Age** | < 60y | 48 | 3.3 | 18.6 | 15.8 | 11.8 | 0.0188 |
|  | 61-80y | 170 | 48.3 | 53.8 | 60.5 | 52.9 |  |
|  | > 80y | 90 | 48.3 | 27.6 | 23.7 | 35.3 |  |
| **Gender** | F | 163 | 73.8 | 42.8 | 63.2 | 52.9 | 0.0009 |
|  | M | 145 | 26.2 | 57.2 | 36.8 | 47.1 |  |
| **Site** | Colon | 253 | 98.3 | 75.9 | 84.2 | 64.7 | < 0.0001 |
|  | Rectum | 55 | 1.7 | 24.1 | 15.8 | 35.3 |  |
| **Side** | Left | 172 | 20 | 71 | 50 | 76.5 | < 0.0001 |
|  | Right | 136 | 80 | 29 | 50 | 23.5 |  |
| **Stage<sup>a</sup>** | 1 | 53 | 16.7 | 17.2 | 28.9 | 11.8 | 0.01449 |
|  | 2 | 128 | 55 | 44.8 | 26.3 | 23.5 |  |
|  | 3 | 105 | 26.7 | 29.7 | 42.1 | 47.1 |  |
|  | 4 | 22 | 1.7 | 8.3 | 2.6 | 17.6 |  |

percentages. N, total number of patients per group; *P*-value\* is derived from analysis of
classified tumours only and a *P*-value of < 0.05 is considered significant; y, years; F, female;
M, male; <sup>a</sup> Post-operative Tumour-Node-Metastasis staging.

### Survival analysis

The median follow-up period was 50 months (0⋅3–172 months) with a median survival of 82 months (95% CI 71⋅8 – 110⋅5). Survival curves and proportions at 5 and 10 years are shown in Figure 1. Both progression-free survival (PFS, *P* = 0⋅039) and overall survival (OS, *P* = 0⋅036) were associated with CSM subtype in the classified samples. The associations were largely due to the difference between CMS subtype 4 and the other classes; the hazard ratios for CMS4 relative to all other classified samples were 2⋅28 (95% CI 1⋅28 – 4⋅05, *P* = 0⋅005) and 2⋅29 (95% CI 1⋅26 – 4⋅18, *P* = 0⋅007) for PFS and OS, respectively. However, after adjusting for age and sex, there was no significant association between CMS stage and OS (*P* = 0⋅11) or PFS (*P* = 0⋅12).

**Figure 1.**
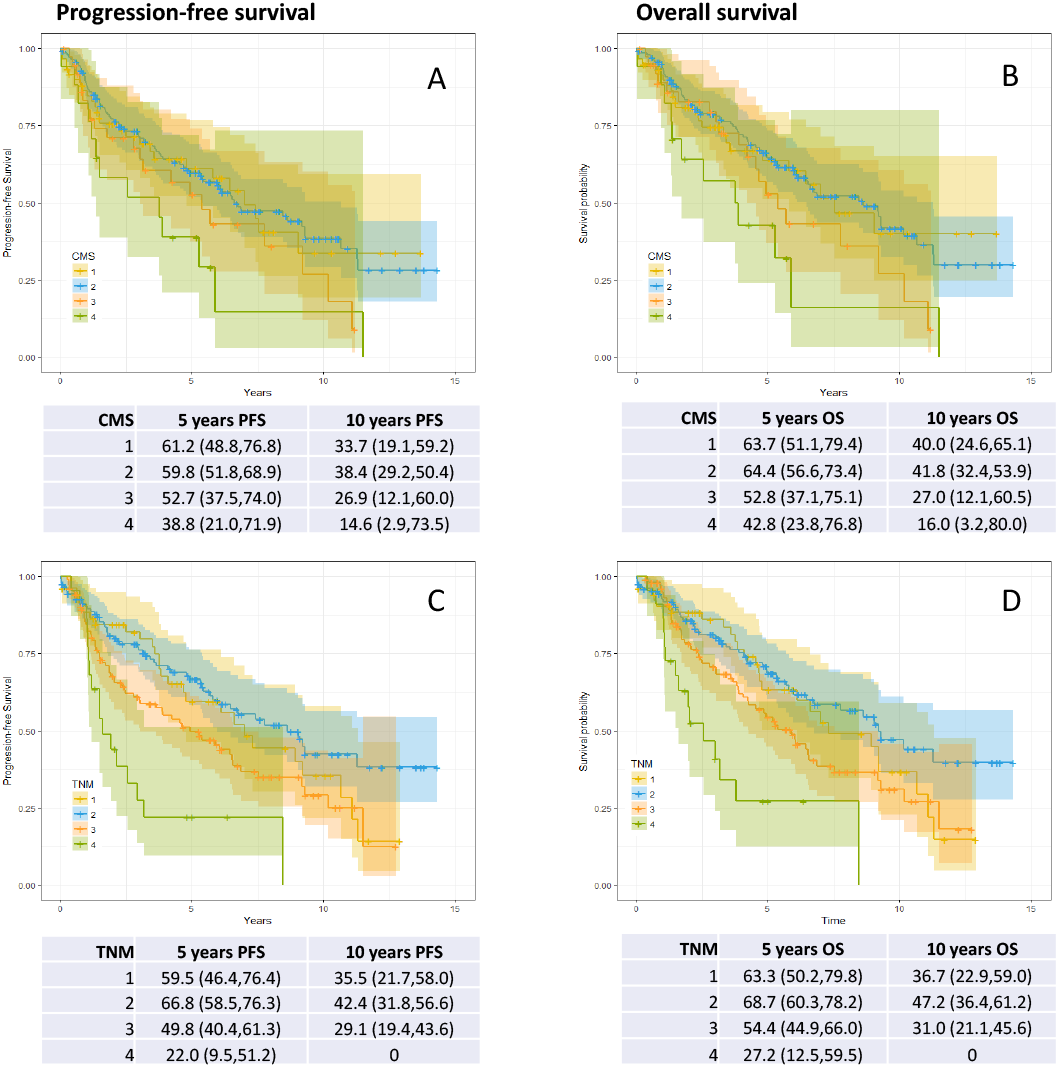
(A) Progression-free and (B) overall survival by consensus molecular subtype (CMS), and (C) progression-free and (D) overall survival by TNM stage. Kaplan-Meier survival curves with estimates and 95% confidence intervals for survival probabilities at 5 and 10 years.

10-year overall survival based on TNM staging showed that, when adjusted for age and gender, Stage 1 and 2 show little difference in survival outcome. However, there is some evidence that Stage 3 is associated with increased mortality, while it is quite clear that Stage 4 is associated with increased mortality (OR = 2⋅8, 95% CI 1⋅6 – 5⋅0, *P* < 0⋅0005).

Considering all samples, older and male patients were at greater risk of poorer outcomes from CRC (Table 4). Cancers that were rectal, had lymph-node involvement, local recurrence or post-operative metastases posed significantly greater risk, however side did not significantly affect risk. Adjusting for all other covariates showed that there was independent risk associated with lymph-node involvement, local recurrence and post-operative metastases, but not rectal cancers. Including both TNM stage and CMS in models of survival analysis shows that TNM stage significantly explains mortality independently of age and gender, whereas CMS subtype does not. From this we conclude that stratification using CMS does not perform as well as TNM staging as an independent prognostic indicator in our cohort.

**Table 4.** Hazard ratios for risk factors associated with mortality in colorectal cancer

|  |  | HR (95% CI) | *P* | *P*<sub>Age,Sex</sub> | *P*<sub>All</sub> |
| --- | --- | --- | --- | --- | --- |
| Age (years) | 60–80 | 2.3 (1.2,4.2) | 0.0002 |  | 0.0002 |
|  | 80+ | 3.4 (1.7,6.5) |  |  |  |
| Male |  | 1.4 (1.0,1.9) | 0.0640 |  | 0.0175 |
| Rectal |  | 1.5 (1.0,2.1) | 0.0511 | 0.1341 | 0.1341 |
| Right side |  | 0.9 (0.7,1.3) | 0.5930 | 0.3881 | 0.7978 |
| Lymph node involvement |  | 1.5 (1.1,2.1) | 0.0139 | 0.0046 | 0.0076 |
| Local recurrence |  | 2.2 (1.2,4.1) | 0.0187 | 0.0120 | 0.0193 |
| Post-operative metastasis |  | 3.5 (2.4,5.0) | <0.0001 | <0.0001 | <0.0001 |
| TNM Stage | 2 | 0.8 (0.5,1.3) | 0.0004 | 0.0000 | 0.0015 |
|  | 3 | 1.3 (0.8,2.0) |  |  |  |
|  | 4 | 3.0 (1.6,5.7) |  |  |  |
| CMS | 2 | 1.0 (0.6,1.5) | 0.0949 | 0.2619 | 0.4237 |
|  | 3 | 1.4 (0.8,2.6) |  |  |  |
|  | 4 | 2.2 (1.1,4.5) |  |  |  |
|  | Unclassified | 1.2 (0.7,2.1) |  |  |  |

Hazard ratio (HR) with 95% confidence interval (CI) and *P*-value from analysis of deviance from univariate models. $P_{\text{Age,Sex}}$ , *P*-value from analysis of deviance for Cox proportional hazards model with covariates for age and gender; $P_{\text{All}}$ , *P*-value from analysis of deviance on Cox proportional hazards model including all covariates; TNM, tumour-node-metastasis; CMS, consensus molecular subtype.

Of the 308 patients, 63 patients had relapse of their disease; either local recurrence or distant metastases or both within the follow-up period. There was no significant difference in the median survival after relapse which was 16⋅5 months, 12⋅4 months, 33⋅9 months and 4⋅6 months for CMS1, CMS2, CMS3 and CMS4 tumours respectively (*P* = 0⋅187)

### Prognostic effect of CMS in CRC stratified by TNM Stage

There were 17 participants with stage 4 cancer classified by CMS and coincidentally 17 CMS4 patients. These numbers were insufficient to draw robust conclusions for stage 4 or CMS4 when cross tabulated, and were omitted from the analysis. Differential survival by stage (1 to 3) and CMS (1 to 3) was identified by analysis of deviance on a Cox proportional hazard model with interaction between TNM stage and CMS (*P* = 0.048). Including covariates for age (dichotomous at 80), sex, tumour site and side increased the significance of this effect (*P* = 0.022). To assess the magnitude of the differences between CMS subtypes within different stage tumours, survival analysis was performed on the data stratified by TNM stage. Median survival times were calculated and Cox proportional hazard models fitted with and without covariates for age, sex, site and side (Table 5). There was a significant difference in survival predicted by CMS for stage 1 tumours. However, this was explained by covariates. For stage 2 tumours, there was a suggestion that CMS subtype 3 has worse survival than CMS1 and 2, which was statistically significant after adjusting for age, sex, site and side. There was no evidence that outcome differed by CMS subtype for stage 3 or 4.

**Table 5.** Survival time by CMS stratified by TNM stage.

|  | CMS1 | CMS2 | CMS3 | *P*-value | *P*adj |
| --- | --- | --- | --- | --- | --- |
| Stage 1 | 39.8 | 127.9 | 110.6 | 0.035 | 0.170 |
| Stage 2 | 108.5 | 121.8 | 50.53 | 0.051 | 0.006 |
| Stage 3 | 90.3 | 72.7 | 64.4 | 0.769 | 0.752 |

Stage, TNM stage; CMS, consensus molecular subtype; *P*adj, *P*-value adjusted for age, sex, site and side

### Differential gene expression and gene-set enrichment analysis in Stage 2 tumours

Differentially expressed genes between Stage 2 patients who died and those who were alive at the end of the follow-up period, were further analysed to identify genes potentially associated with survival in Stage 2 tumours. Differentially up-regulated genes strongly associated with survival in this patient group includes immune-cell related genes, in particular genes coding for B-cell markers, and several known (*LRRC4*, *PKNOX2*, *FEZF2*) and putative tumour suppressor genes (*MTO18B* and *NCAM1*) (Supplementary Table S3). Genes that were significantly up-regulated in patients with poor survival included pro-inflammatory genes (*IL17REL*, *RETNLB*) and genes that have been previously associated with progression and poor outcome in CRC (*ERBB2*, *TBLRXR1*, *TAPBP*, *CPS1*, *AGR2*) (Supplementary Table S4).

In order to compare biologic pathways and processes potentially associated with survival in this subgroup of patients, we used DEGs as input into an assortment of gene ontology tools. Differentially upregulated biologic pathways associated with survival were predominantly immune pathways, including B-cell activation, IL-12 and PD-1 signalling and T-cell activation. In addition, glutamatergic signalling was differentially enriched in Stage 2 patients still alive at the end of follow-up (Supplementary Table S5). GSEA showed an enrichment of pathways involved in metabolic regulation in Stage 2 tumours, which reflects the association of CMS3 with poor survival, and also differential up-regulation of processes involved in protein and nucleic acid synthesis (Supplementary Table S6).

## 4. Discussion

Recent advances in gene expression analysis have culminated in the publication of a Consensus Molecular Subtyping (CMS) system by the Colorectal Cancer Subtyping Consortium (CRCSC) that stratifies CRC into one of four subtypes based on transcriptional profiling. The CRCSC study reported an association between CMS4 and worse patient outcome, and between CMS1 and survival after relapse. Although many subsequent reports mention the prognostic potential of CMS, no study, to date, has validated the prognostic impact of the subtyping system compared to the routinely used staging for primary CRC.

In a large, single-institution cohort of chemotherapy-naïve, surgically treated colorectal cancers, we have shown that traditional TNM staging outperforms molecular subtyping in prognostication of CRC. Post-surgical staging of this cohort was carried out according to UICC guidelines and staging was similar to that expected of a treatment-naïve cohort. Association of stage with clinical variables found few associations beyond the parameters used to carry out staging, namely lymph-node involvement and distant metastases. As previously reported for other cohorts,[18, 19] the association of increasing stage with metastasis was also largely driven by liver metastases.

In addition to histological staging, we carried out consensus molecular subtyping (CMS), based on RNA-sequencing derived gene-expression profiles from tumour tissue. Stratification into CMS yielded similar proportions of CMS1 and CMS3 and unclassified tumours as described by the CRCSC.[8] Our cohort contained a considerably greater proportion of CMS2 tumours at 47%, compared to 37% reported by CRCSC, and fewer CMS4 tumours, 6% compared to 23%. The difference in reported proportions of CMS may be, at least in part, accounted for by the inclusion criteria of surgery with curative intent and the exclusion of patients who received neo-adjuvant chemo-or radiotherapy in this study. A recent report by Trumpi et al reported that neoadjuvant therapy induces a mesenchymal phenotype in residual tumour cells and, as such, may lead to an increase in the reporting of CMS4 subtypes.[20] These criteria may have excluded many advanced-stage tumours, which were shown to be associated with CMS4.[8] Intra-tumoural heterogeneity may also affect the classification of CMS4 tumours, as the EMT-associated genes seen in CMS4 tumours may reflect upregulated genes derived from fibroblast and mesenchymal cells present in the stromal background rather than directly from the tumour itself,[9, 21-23] and several studies have suggested that the location and number of tumour biopsies can undermine the accuracy of CMS;[24–26] a limitation of this study is the use of a single tumour sample to carry out gene-expression profiling.

Stratification into CMS showed similar associations with clinic-pathological variables as previously reported by CRCSC and other studies. CMS1 tumours were associated with right-side, female, node-negative and poorly-differentiated, with a high proportion of mucinous histology and less likely to be seen in younger patients under 60 years of age. CMS2 tumours made up nearly half of our cohort and were predominantly left-sided tumours found in male patients, and showed a negative association with mucinous type. Patients with CMS3 type tumours were associated with a lower TNM stage. CMS4 tumours were associated with younger age, rectal tumours and presented at an advanced TNM stage with lymph node positivity. Indeed, lymph-node positivity is shown to increase through CMS1, 2, 3 to CMS4.

The established association between increasing tumour stage and poorer outcome was recapitulated in our cohort, in terms of progression-free and overall survival. While post-surgical staging is the mainstay of prognostication in most clinical centres, the potential for refining prognostication using molecular features has been widely investigated, and the combined use of different clinical and molecular markers have shown links with prognosis in CRC, e.g. while *BRAF* mutations have been associated with poorer outcome,[27] the effects of these mutations may be mitigated in MSI tumours.[28, 29] *KRAS* mutations are also associated with a poorer outcome, but this association is stronger in distal compared to proximal tumours.[30]

The original study by Guinney et al first describing consensus molecular subtyping showed an association between CMS4 and poor overall survival, and between CMS1 and survival after relapse.[8] Several studies have incorporated CMS with other molecular features to in order to refine prognostic groups, and have described poorer outcomes in *BRAF*-mutated CMS1 MSS tumours, and *KRAS-*mutated CMS2/3 MSS tumours[31], and favourable outcomes in CMS1 MSI tumours.[32] Although many subsequent publications have emphasised the prognostic importance of CMS, the utility of CMS as a stand-alone prognostic tool in the clinical setting has not been investigated in an independent cohort. Survival analysis showed an association between CMS and both progression-free and overall survival in our cohort, and this was largely due to the difference between CMS4 and the other CMS classes. However, after adjusting for age and sex, CMS4 was not an independent prognostic marker for survival in this study. Including both TNM stage and CMS in models of overall survival shows that TNM significantly explains mortality independently of age and gender, whereas CMS does not. A potential limitation of the study is the relatively low numbers of CMS4 tumours, as discussed above, and that almost half of the tumours in our cohort are CMS2, and this imbalance may affect the power of our study to detect effects specific to CMS1, 3 and 4.

Clinical management of CRC is usually based on histological staging, with stage 1 tumours conservatively managed with surgery and tumours with nodal or distant metastases (Stage 3 and 4) usually treated with adjuvant chemotherapy. Stage 2 tumours remain a conundrum in terms of prognostication, as approximately 20% of patients with Stage 2 CRC die from the disease[33]. Various factors including acute presentation with obstruction and perforation, histological factors such as perineural and perivascular invasion, as well as high grade, have been used as markers of poor prognosis, and as such indicators for adjunctive postoperative chemotherapy. Further stratification using molecular markers, such as *BRAF* and *KRAS* mutations, and MSI status[34] have been investigated with regard to their prognostic potential in this tumour group, but have not widely adopted to direct clinical management. Molecular subtyping is a cornerstone of precision medicine in cancer treatment, and the mutation status of genes in the EGFR pathways, including *RAS* genes, *PIK3CA*, *PTEN* and *BRAF* have been shown to predict response to EGFR blockade therapy in CRC.[37] MSI status and the effect of the tumour microenvironment, in particular the amount and type of tumour infiltrating lymphocytes, have more recently been proposed as predictors of response to immunotherapy.[38] To date, although CMS1 tumours encompass a large proportion of MSI positive CRC, no targeted treatment options based solely on CMS have been proposed. In the context of metastatic CRC, CMS appears to associate with survival in clinical trials of patients with wild-type *KRAS* tumours, treated with anti-EGFR or VEGF inhibitors[39, 40]. However, in primary CRC and outside the clinical trials setting, improved stratification of CRC had not yet been demonstrated using CMS. In this study, we have observed that subtyping of TNM-stratified tumours into CMS could improve prognostication for Stage 2 CRC; tumours that were CMS3 subtype had significantly lower overall survival compared to other molecular subtypes. This demonstrates, for the first time, the potential utility of CMS in improving prognostication of CRC in combination with existing methods.

Differential gene expression between Stage 2 patients who died and those who were alive at the end of the follow-up period, identified significant up-regulation of immune-related genes and biologic processes, and tumour-suppressor genes, associated with survival. The importance of the immune microenvironment in tumour progression has been demonstrated in solid tumours, and has been linked to outcome in CRC. Our findings suggest that immune signatures may identify Stage 2 patients with good prognosis, for whom surgery alone may suffice, and conversely a CMS3 signature may identify patients who would benefit from adjuvant chemotherapy/increased surveillance. Further evaluations of the genetic signatures identified in this study, and prospective validation using a more clinically accessible platform e.g. gene panel test, will be necessary to confirm these findings.

CMS currently represents the best description of tumour heterogeneity in colorectal cancer at the level of gene expression, and shows promise for future advancement of precision medicine. Our findings also suggest its use in refining prognostication in the clinically heterogenous Stage 2 colorectal cancer.

### Ethics approval and consent to participate

Ethical approval was granted by the University of Otago Human Ethics Committee (ethics approval number: H16/037). Study participants gave informed written consent, and the study was performed in accordance with the Declaration of Helsinki.

### Availability of data and materials

All sequencing data will be archived in Sequence Read Archive upon acceptance of the manuscript for publication. Additional information (bioinformatics code and limited patient metadata) will be provided upon reasonable request to the corresponding author.

## Funding Sources

Maurice and Phyllis Paykel Trust.

Gut Cancer Foundation (NZ), with support from the Hugh Green Foundation.

Colorectal Surgical Society of Australia and New Zealand (CSSANZ).

The Health Research Council of New Zealand.

The funding bodies had no role in the design of the study or collection, analysis, and interpretation of data or in writing the manuscript.

## Authorship

R.P. carried out sequencing preparation of tumour samples, and was a major contributor to study design and manuscript writing. S.S. carried out bioinformatics analysis and preparation of figures and contributed to manuscript writing. Y.L was involved in collated patient data and statistical analysis. J.P was involved in study design, bioinformatics and data analysis, and manuscript preparation. F.F. was involved in study design and clinical aspects of the study. All authors read and approved the final manuscript.

## Acknowledgement

The authors would like to thank Helen Morrin at the Cancer Society Tissue Bank, Christchurch, and the patients involved for generously participating in this study.

